# A Comprehensive Method on Black-legged Tick Larvae and Nymph Feeding on Mice to Study Lyme Disease Transmission and Acquisition

**DOI:** 10.1101/2024.11.20.624518

**Authors:** Aaron Scholl, Bingjie Li, Sandip De

**Affiliations:** Tumor Vaccines and Biotechnology Branch, Division of Cellular Therapy 2, Office of Cellular Therapy and Human Tissue, Center for Biologics Evaluation and Research, U.S. Food and Drug Administration, Silver Spring, Maryland, USA

**Keywords:** tick, Lyme, *Borrelia burgdorferi*, mice, feeding, transmission, acquisition

## Abstract

Tick-borne diseases are a growing public health concern in the United States, with cases rising steadily each year. Lyme borreliosis, or Lyme disease, remains the most prevalent, affecting approximately 476,000 individuals annually. Human-driven changes in climate and ecosystems have expanded the habitat of pathogen-carrying ticks, facilitating the spread of these infections. Additionally, increased instances of tick-borne diseases transmission through human tissues have been reported. Despite ongoing efforts to manage these infections, their incidence continues to rise. To develop effective control measures against these diseases and prevent the transmission of tick-borne infections through human and animal tissues, it is very important to develop detection assays and understand the transmission mechanisms of tick-borne infections. In this study, we provide detailed descriptions and visual references for larval and nymphal tick feeding on mice, focusing on the transmission and acquisition of *Borrelia burgdorferi* (sensu stricto). These methodologies can be applied to study other tick-borne diseases, tick vectorial capacity, and tick biology, aiding in the development of detection strategies to combat these infections.

## Introduction

Ticks are becoming more prevalent worldwide as vectors of diseases affecting humans, livestock, and companion animals. This expansion has been exacerbated due to changes in climate that are becoming more hospitable to ticks year-round. In addition, deforestation, reforestation, and development over recent decades are believed to play a key role in the spread of ticks. There are more than 900 tick species that feed on mammals, birds, and reptiles. In the United States, at least 9 different tick species are known to spread diseases to humans (1). The deer tick, or black-legged tick (Ixodes scapularis), is the most significant tick species responsible for transmitting at least seven different pathogens to humans, including *Borrelia burgdorferi*, which causes Lyme disease (2). This tick species is found throughout much of the eastern and central regions of North America, including the United States, Canada, and Mexico.

Deer tick has three different developmental stages-larvae, nymph, and adult, they are most active in the spring and fall but can be active year-round in warmer climates where temperatures remain above 4°C (2, 3). They are also known to feed on a wide variety of mammals, including deer, rodents, and humans. Ticks spread multiple pathogens to humans, including anaplasmosis, babesiosis, and Lyme disease (4, 5). While some pathogens are native to ticks, others, such as *Borrelia burgdorferi*, are acquired via ticks feeding on infected hosts (6). The CDC estimates that roughly 476,000 patients were diagnosed and treated for Lyme disease annually in the United States between 2010 to 2018, up from an estimated 329,000 patients between 2005 and 2010 (7, 8).

The rising incidence of Lyme infection raises concerns about increased disease transmission through tissue transplantation. For instance, there have been reports of active Lyme disease being transmitted following penetrating keratoplasty (9). In addition, transplant and transfusion-transmitted infections have been documented for other tick-borne pathogens, including *Babesia, Anaplasma phagocytophilum, Rickettsia* and *Ehrlichia* (10-20). Effective control of tick-borne infections requires a deeper understanding of transmission dynamics and the development of improved detection assays. Here, we have provided a detailed methodology for studying *Borrelia* transmission, acquisition, and detection by feeding deer ticks on mice. This methodology can be also adapted to study the dynamics of other tick-borne infections and to develop improved detection assays in the laboratory.

## Materials and Methods

Ethics Declaration: The study was conducted following approval by the Institutional Animal Care and Use Committee (IACUC) at CBER/FDA, (Protocol Number #08-2022). The study adhered to the animal research ethics committee guidelines of CBER/FDA.

### Personal Protective Equipment

The proper use of personal protective equipment (PPE) is mandatory when working with ticks. PPE requirements vary by institution and country. Full coverage PPE is highly recommended during the entire feeding procedure, and should consist of a hair net, surgical mask, coveralls/white gown, white/light colored gloves, and shoe covers. A depiction of full coverage PPE is shown in **Figure 1A**. PPE should be white or light-colored to aid in the visualization of ticks. Proper coverage of PPE provides basic protection as a physical barrier, preventing any ticks from crawling inside and attaching to the skin of research or animal facility staff. A hair net is worn to cover the entire crown of the head and extends below the ears as demonstrated in **Figure 1B**. Adhesive white mats (Fisher Scientific, Cat# 19-181-510) were used as another method of tick-containment and were placed under any container and on any surface where ticks were expected to be handled. The adhesive white mat aids in sticking and visualization in the unlikely event that a tick climbs out, falls, or drops during handling.

**Figure 1.**
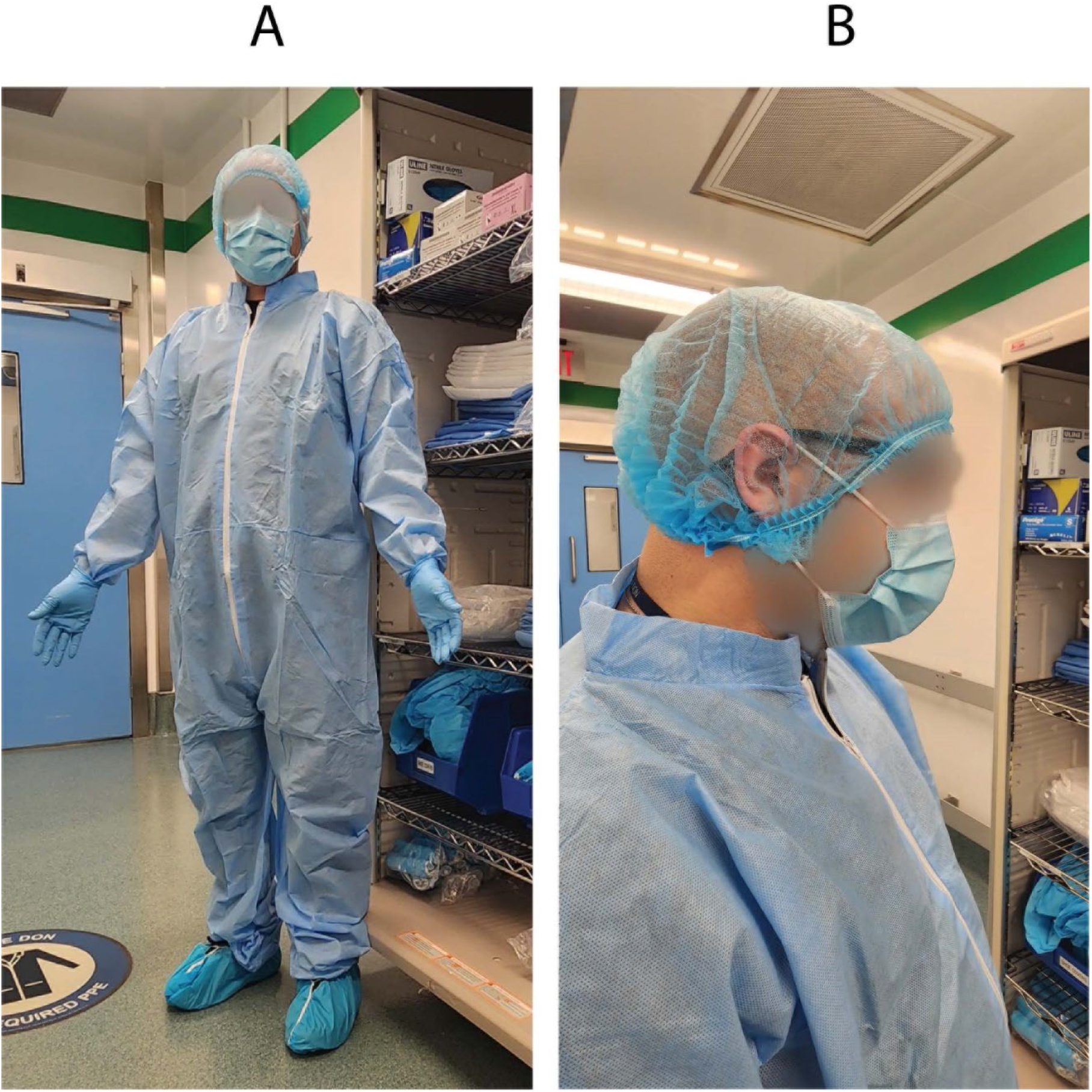
Personal Protective Equipment. (A) Full coverage PPE consists of a hair net, surgical mask, coveralls/white gown, white/light colored gloves, and shoe covers. Gloves are then donned over the hand and sleeve to reduce skin contact areas. (B) The hairnet is pulled below the ears to secure it to the head.

### Tick Storage

Black legged tick larvae and nymphs were purchased from tick rearing facility at Oklahoma State University. Prior to feeding the larvae and nymphs, the ticks were stored on a rack inside of a transparent closed chamber (Secador 2.0 vertical desiccator cabinet, SP Bel-Art) that was placed on an adhesive white color mat stored within an incubator (Fisherbrand Isotemp incubator) set to 25°C (**Figure 2A, B**). Distilled water in an open container was placed on the bottom rack of the chamber to provide and maintain 90% humidity within the chamber (**Figure 2A, B**). The ticks were stored within a 10×75mm polystyrene test tube (Axiom Medical Supplies M-1035956-3123) with fine nylon mesh (Voile White Decorator fabric) placed between the tube and flange 12 mm polyethylene plug top (Universal Medical Inc. GS-118139C) **(Figure 2C)**. Several holes were poked into the polyethylene plug top with a hypodermic needle to allow for airflow (**Figure 2D**). Double-sided tape (Scotch) was placed around incubator door openings to further prevent any ticks from escaping (**Figure 2E**). Two LED light strips were cut to 12 LEDs each, connected to the same power supply, and installed on opposing sides of the internal top of the incubator (**Figure 2F**). The connected LEDs were run through a programmable timer with a 12:12 light:dark cycle (**Figure 2G**) (Fisherbrand Traceable Digital Outlet Controller, Cat# 06-662-24).

**Figure 2.**
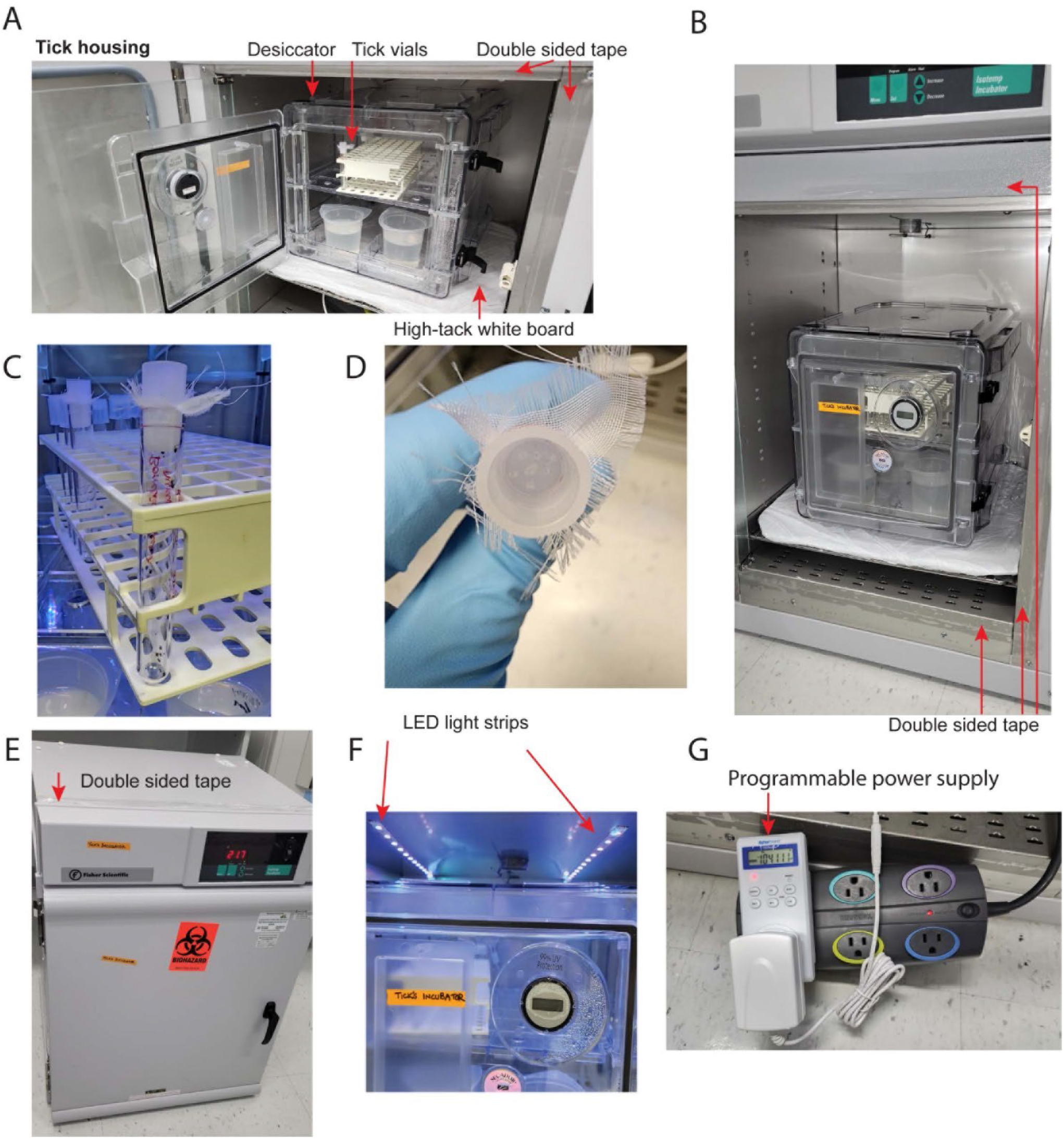
Incubator Setup. (A) Tick housing contains a desiccator chamber with open containers of distilled water for humidity, a rack, and vials for housing ticks. The surrounding edges of the incubator covered in double sided tape to provide a barrier to escaping ticks. The desiccator chamber placed on a high tack white mat as preventative barrier to tick escape. (B) Double sided tape is adhered around the external portion of all chamber walls. (C) A polystyrene vial containing ticks, with fine nylon mesh placed between the tube and flange 12 mm polyethylene plug top. (D) Several holes poked into a polyethylene plug top via a hypodermic needle to allow for airflow. (E) Double sided tape is adhered to external surfaces of incubator. (F) Two pairs of 12 LED lights per strip were adhered to the top of the interior chamber wall. (G) The LED strips were plugged into a programmable power supply. A power strip was included in the series for space concerns, as the chamber’s internal power outlet is directly behind the shelf.

### Infecting Mice with *Borrelia burgdorferi*

To study non-native pathogen transmission, ticks must be infected with the pathogens, and for *Borrelia burgdorferi*, mice need to be infected for the tick nymphs/larvae to feed on them. For our research, we purchased *Borrelia burgdorferi* from BEI resources (NR-13251, strain B31). Before infecting mice, *Borrelia burgdorferi* was cultured in Barbour-Stoenner-Kelly (BSK) medium in the laboratory (Sigma-Aldrich, B8291) (21). Post-culturing, the bacterial concentration was determined with a Petroff-Hausser counting chamber and a dark-field microscope. Exponentially growing bacterial stock was diluted to 10^6^/ml in BSK broth and 100μl (10^5^ total *Borrelia*) was injected per C3H mouse intradermally. A visible bubble should form at the injection site when performed properly. Ticks were infected with B. burgdorferi by placing and feeding them on *B. burgdorferi*-infected mice 15 days after injection as described below.

### Preparing the Cages for Tick Feeding on Mice

In our protocol, ticks were fed on C3H mice, as C3H develops Lyme arthritis (22). Mice are housed in standard plastic mouse cages (11-1/2” long x 7-1/2” wide x 5” deep) with stainless steel wire flooring during tick feeding. Mice are provided with standard food and water during tick feeding. A clear cylinder was added to each cage as enrichment and as a resting place from the wire flooring. Roughly ¼ inch of water was added to the bottom of the mouse cage prior to adding the wire (**Figure 3A**). Ticks that have fed and detached, drop off the mouse and float in the water below. Mice may attempt to consume fed ticks that climb the walls. The top ½ inch of the interior of the mouse cages were smeared with petroleum jelly, which acts as a second barrier for any ticks that have fed, detached, and climbed the walls of the cage (**Figure 3A**). Each prepared mouse cage was placed into a larger guinea pig cage (19” long x 10-1/2” wide x 8” deep) that was also prepared with 1 inch of water and the top ½ inch of the interior smeared with petroleum jelly (**Figure 3B**). The mouse cage inside of the guinea pig cage was then kept on a white sticky mat on a metal shelf.

**Figure 3.**
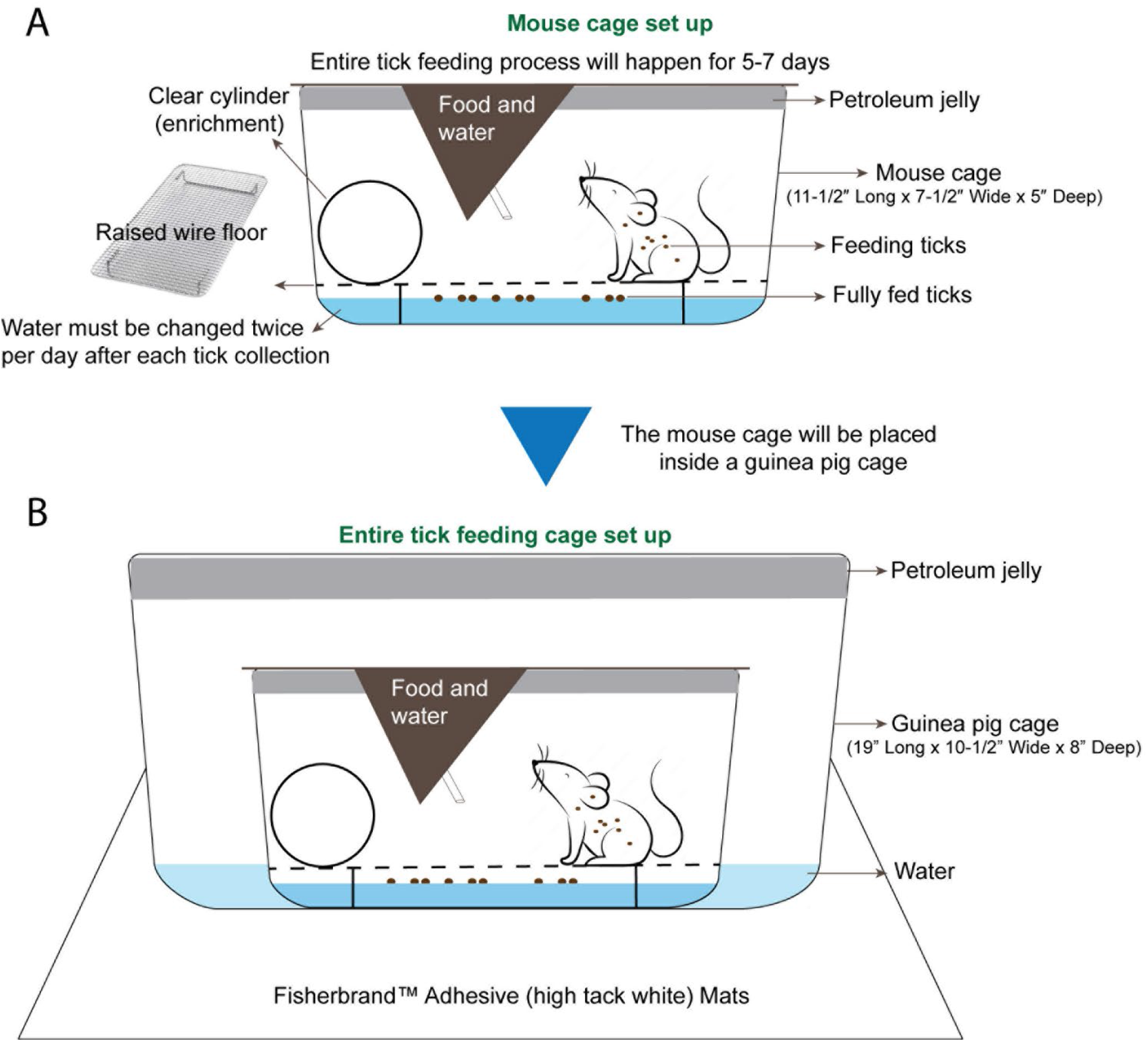
Mouse Cage and Tick Feeding Setup. (A) The mouse cage (11-1/2” Long x 7-1/2” Wide x 5” Deep) is prepared with petroleum jelly around the top ½” of the cage. Roughly ½” of water is added and a raised wire floor is placed inside the cage. A clear cylinder is added for enrichment and as a resting place from the wire floor. (B) A guinea pig cage is prepared with petroleum jelly around the top 1/2” of the cage, and roughly 1/2” of water is added to the guinea pig cage. The prepared mouse cage is placed inside of the guinea pig cage and the whole setup is placed on white adhesive mats on a shelf of the animal room.

A 10% bleach solution is prepared fresh daily to sanitize any collected waste liquids and to kill any uncollected ticks and tick-borne pathogens that were present in the wastewater. A 5-gallon jerrican was designated for 10% bleach disposal. Waste liquids that were treated with bleach for 24 hours were discarded.

To feed the ticks, mice were anesthetized (100mg/kg (ketamine) + 10mg/kg (xylazine), 0.1ml/100g intraperitoneally) and placed next to each other in a clean and empty mouse cage with petroleum jelly smeared on the top interior (**Figure 4A**). A heating pad was placed under the cage for supplemental heat while the mice were anesthetized. Ophthalmic ointment was applied to the eyes of each mouse to prevent drying while anesthetized, as the eyes remained open during anesthetization.

**Figure 4.**
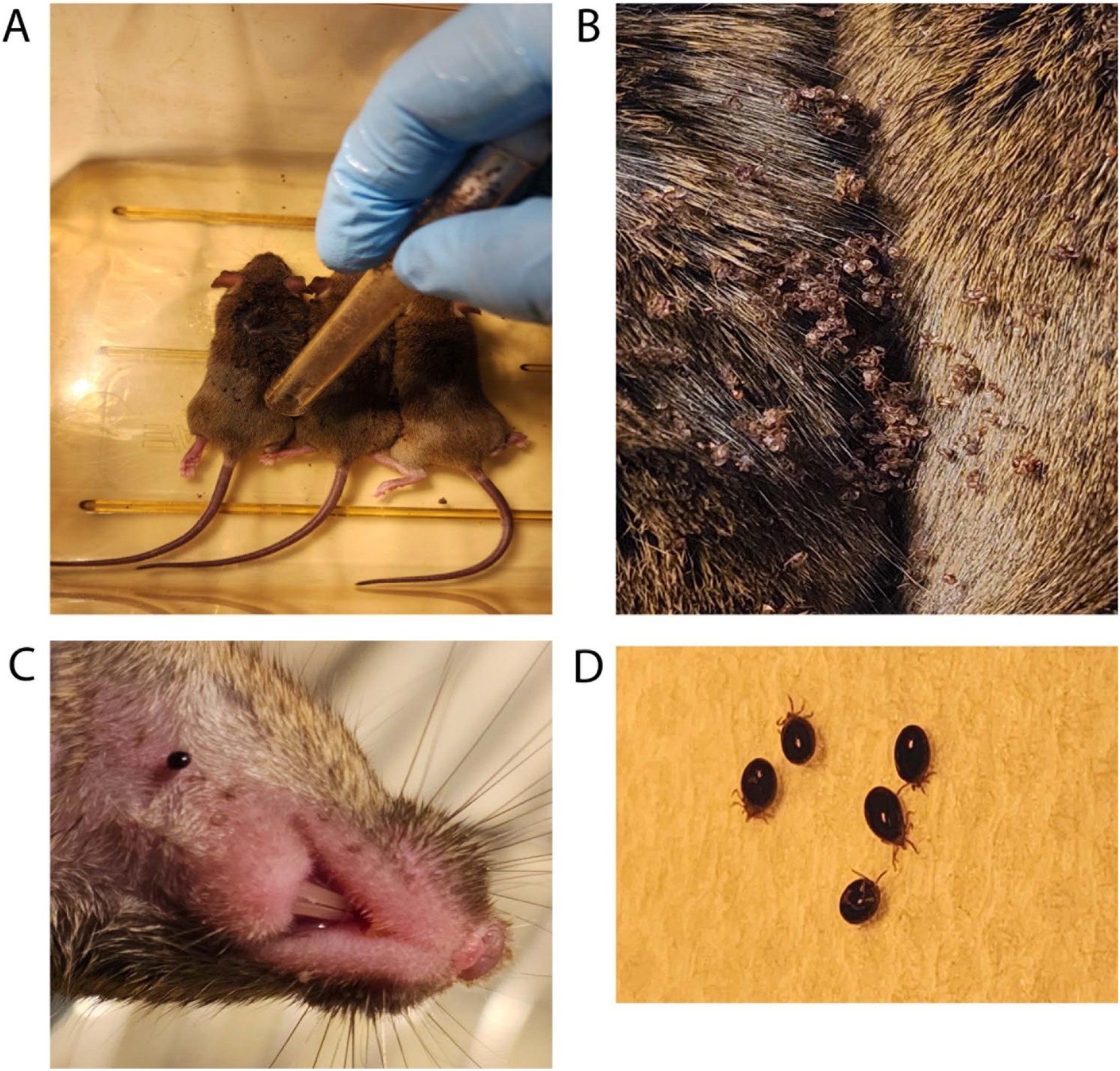
Larval ticks feeding on mice. (A) Tick larvae are introduced from a storage tube that has been smeared on the top half with petroleum jelly. (B) Tick larvae are allowed to disperse while mice are recovering from anesthesia. (C) A closeup view of a larval tick feeding on host mouse neck. (D) Shown are larval ticks that have become fully engorged, dropped off into the wastewater, and been rinsed with clean water.

### Larval Tick Feeding

1. Following anesthetization of the mice, the top half of a tube containing one larval batch was smeared with petroleum jelly. Larvae were tapped down by placing an index finger over the lid and firmly tapping the bottom of the tube on the table. After confirming the absence of larvae on the lid, the tube was moved next to the mice and the lid was opened near the back and neck of the anesthetized mice.
2. After releasing larvae on mice (**Figure 4A**), the lid and tube were quickly submerged into a 10% bleach solution to kill any remaining larvae.
3. The anesthetized mice were directly observed for 30-90 minutes and allowed to awake from anesthesia. This period of inactivity also allows the larvae time to attach to the mouse (**Figure 4B**).
4. After recovering from anesthesia, the mice were transferred into the individually prepared cages where ticks were observed feeding on mice (**Figure 4C**).
5. Over the course of seven days, fed larvae were collected from the wastewater and cages with flat, white-bristled size 24 and size 2 paint brushes, respectively. This collection was performed in the morning and evening on each of the seven days, roughly eight to ten hours apart. The mice were transferred to a fresh tick feeding cage while the fed larvae were collected from the wastewater and cage (**Figure 4D**).
6. Fifty fed larvae were then transferred into each round-bottom 10×75mm polystyrene test tubes with fine nylon mesh placed between the tube and flange plug top with airholes poked in it. To avoid larval death from mold growth in the collection tubes from exposure to wastewater, the fed larvae were transferred to new storage tubes after a week.
7. The tube of fed larvae was placed on a rack inside the storage container detailed above. Fed larvae can be processed for DNA, RNA, or protein, or allowed to molt into nymphs, depending on experimental needs.

For investigating *Borrelia burgdorferi* transmission to mice and detection thereof, 3-5 *Borrelia*-infected nymphs should be placed on naïve mice. For *Borrelia* acquisition and xenodiagnostic studies, one should put not more than 25-30 nymphs on infected mice (as described below).

### Nymphal Tick Feeding

8 Following anesthetization of the mice, to introduce the tick nymphs, the top half of the tube containing nymphs was smeared with petroleum jelly. The nymphs were tapped down by placing an index finger over the lid and firmly tapping the tube on the table. After confirming that no nymphs were on the lid, the lid and mesh were removed carefully and Dumont tweezers style 5a (EMS #72720-D) were used to individually select and hold the nymphs by their posterior legs. A fine-bristled paint brush can also be used to place the tick nymphs onto the mice.
9 During handling, caution was taken to avoid handling the ticks by their first pair of legs. Haller’s organs are sensory organs present on the terminal portion of the first legs that, if damaged, can prevent the tick from finding and attaching to a host skin.
10 Nymphs were placed onto the back, neck, and ear of the anesthetized adult mice. Ticks were placed under the fur (**Figure 5A, B**). A maximum of 30 nymphs were placed per adult mouse. Empty tubes were placed into a 10% bleach solution for 24 hours before discarding. The number of nymphs placed on each mouse was recorded.
11 The anesthetized mice were directly observed and allowed to awake from anesthesia as described above. This period of inactivity allows the nymphs time to attach to the mouse.
12 After recovering from anesthesia, each mouse was transferred to their individually prepared cages.
13 Nymphs were collected from the mice at experimentally determined intervals after placement or collected as fully fed ticks over the course of 7 days. The partially fed nymphs were picked from the skin of the mice using Dumont style 5a tweezers and placed into microcentrifuge tubes for transport. The partially fed ticks were immediately dissected and processed. Partially fed nymphs can be dissected for specific tissues to extract DNA/RNA/protein.
14 Tubes of fed nymphs were placed on a rack inside of a humidified chamber in a 25°C for storage.

**Figure 5.**
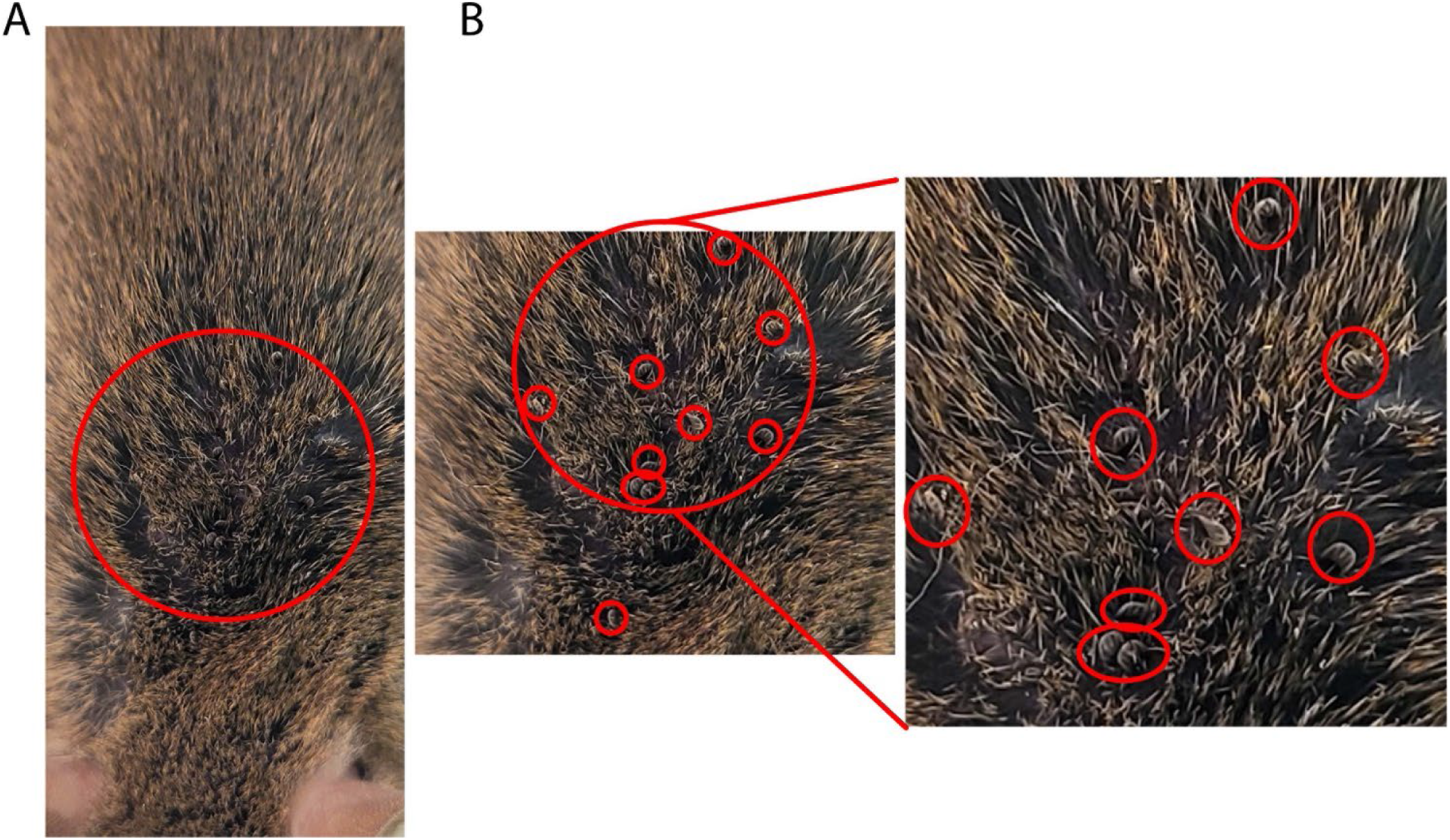
Nymph ticks feeding on mice. (A) Introduction of tick nymphs by placing into fur with tweezers or brush. (B) Multiple tick nymphs moving into mouse fur. Zoomed view with red circles indicating tick location amongst mouse fur.

During the entire feeding process, the tick-infested mouse was transferred to a freshly prepared tick feeding cage every morning and evening, with the wastewater having been thoroughly checked for ticks. Wastewater was treated with bleach as described above. During the cage transfer of mice, cages should be kept on a white sticky mat as described above. By Day 3 of tick feeding, 10-50% of ticks drop off after becoming fully engorged. By Day 4, 95-98% of ticks dropped off the mice. Day 5 contained the lowest yield of fully fed ticks, with the subsequent days yielding no additional ticks. Following tick feeding, the mice were euthanized via asphyxiation and cervical dislocation. Mouse blood and tissues can be harvested at this step to investigate various avenues of tick-host interaction.

### Dissecting Ticks to Collect Specific Organs for Staining or RNA Extraction

The tick nymph was placed into a drop of PBS on a sterile microscope slide (**Figure 6A**). Using an Olympus SZ61 dissecting microscope to visualize, the legs were each removed by holding the tick with tweezers and cutting off the legs with vannas scissors (World Precision Instrument 501839). Once the legs were removed, the tick is rendered immobile and easier to dissect. While holding the chelicerae with tweezers, a flat cut was made with vannas scissors on the most posterior portion of the tick (**Figure 6B**). Two opposing diagonal cuts at roughly 45 degrees and 135 degrees were made connecting with the previous cut, cutting just laterally to the spiracular plates of the trachea (**Figure 6B**). Using the newly created opening (**Figure 6C**), the interior of the tick is exposed by peeling the dorsal scutum from the ventral exoskeleton. The scutum was torn away from the ventral exoskeleton, as the ventral side contained all major anchor points for internal tissues (**Figure 6D**). Care was taken during scutum removal to not disrupt the head of the tick and the scutum was discarded (**Figure 6D**). Both the salivary gland and midgut were located and the tracheal tissue surrounding these organs was removed by gently tearing them away. The midgut was removed first, as it sits on top of the salivary glands (**Figure 6E**). Using two pairs of tweezers, each portion of the midgut was carefully separated from surrounding tissue to prevent excess tearing, contamination, and leakage. The midgut was placed onto a slide with 1 drop of 1x PBS and fixed with acetone for immunofluorescence assays or into a microcentrifuge tube with 100μl TRIzol for RNA extraction.

**Figure 6.**
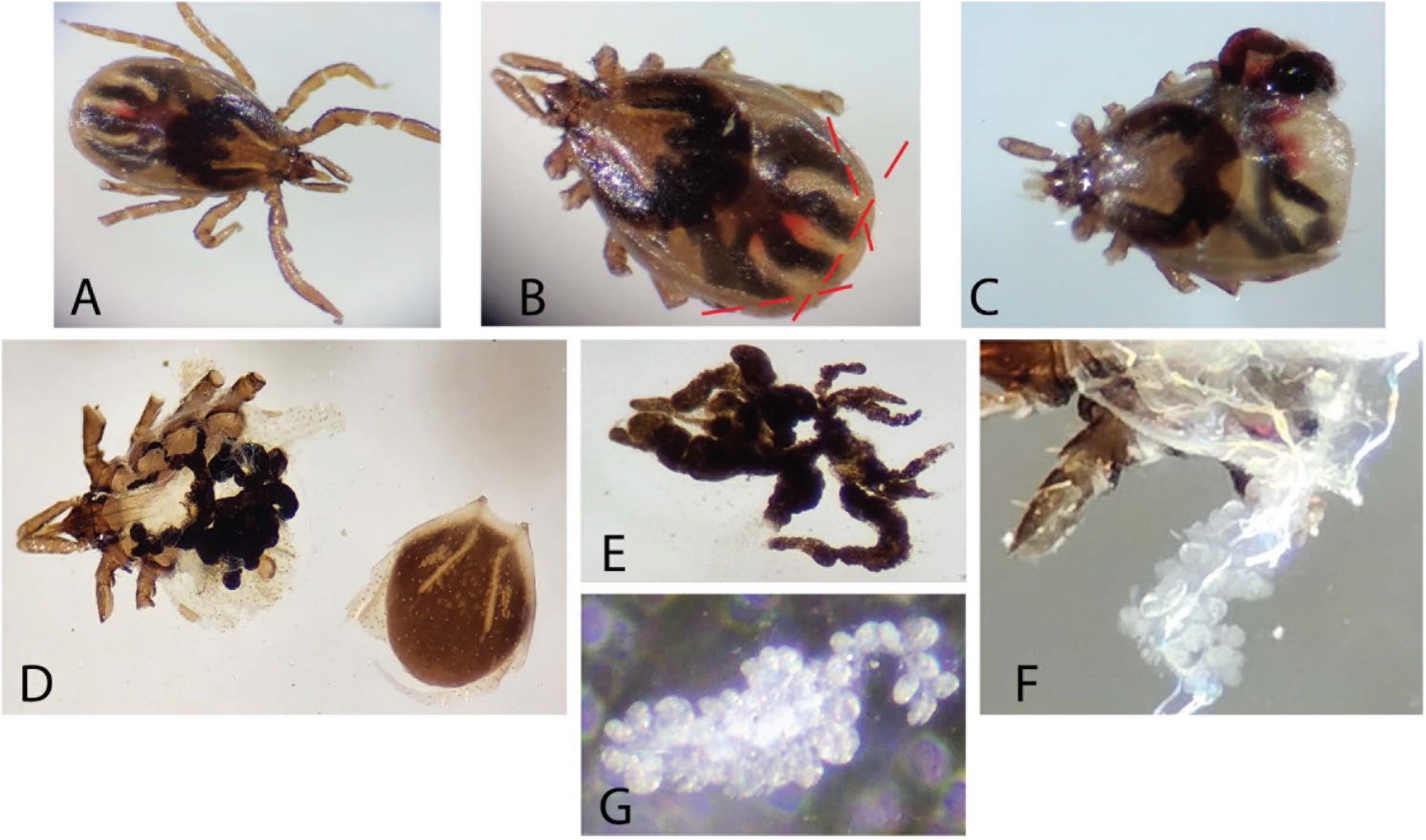
Tick dissection. A 24-hour fed nymph is depicted in A-C, while an unfed nymph is depicted in (D-G). (A) An undissected 24-hour fed nymph is depicted. (B) The legs were carefully cut away to prevent tick mobility and subsequent cuts are indicated as dashed lines. (C) Cuts were made with vannas scissors at the posterior along with intersecting 45- and 135-degree cuts to expose internal organs. (D) The scutum was detached by careful tearing with fine-nosed tweezers. (E) The midgut was isolated from the tick. (F) The salivary glands appeared as grape-like clusters after the midgut has been removed. (G) The salivary glands were carefully isolated from the surrounding tracheal tissues.

Following midgut removal, both salivary glands, which appears as spherical clusters (**Figure 6F**), were dissected from their respective sides by tearing the anterior lobular salivary duct nearest to the head (**Figure 6G**), then placed onto a slide with PBS and fixed with acetone for immunofluorescence assays or into a microcentrifuge tube with 100μl TRIzol for RNA extraction.

### Tissue Fixation and Immunofluorescence Assay of *Borrelia burgdorferi*-infected samples

Tick guts and salivary glands were washed with 3 drops of PBS after dissection. Post-washing, acetone was added and treated for 10 minutes to fix and permeabilize the tissues. After fixation, tissues were blocked using PBS-T (0.05% Tween-20 in PBS) supplemented with 5% normal goat serum for 30 minutes at room-temperature (RT). The blocking buffer was removed, and samples were incubated with primary antibody [Anti-*Borrelia* (Abcam, ab69219) diluted in fresh blocking buffer overnight at 4°C (primary incubation can also be done for 1 hour at RT). Samples were washed three times in PBS-T for 5-10 minutes each at RT before being incubated with secondary antibody [Goat anti-Rabbit Alexa Fluor® 488 (ab150077) for 1 hour at RT. During and following secondary antibody incubation, samples were protected from light by placing them in a foil-wrapped container during incubations. Samples were then washed three times in PBS-T for 5-10 minutes each at RT. Following washes, the tissues were transferred onto Poly-Lysine coated slides, and a drop of anti-fade mounting media with DAPI was added to the samples before applying a cover slip. Samples were incubated with mounting media for at least 20 minutes prior to imaging. Positive *Borrelia* infection results of tick gut tissues are depicted (**Figure 7A, B**). Western blots of uninfected mammalian cell lysate are compared to cultured *Borrelia* lysate probed with *Borrelia*-infected mouse serum (**Figure 7C**) or an anti-*Borrelia* commercially produced antibody (**Figure 7D**) indicate positive *Borrelia* signals in the infected mouse.

**Figure 7.**
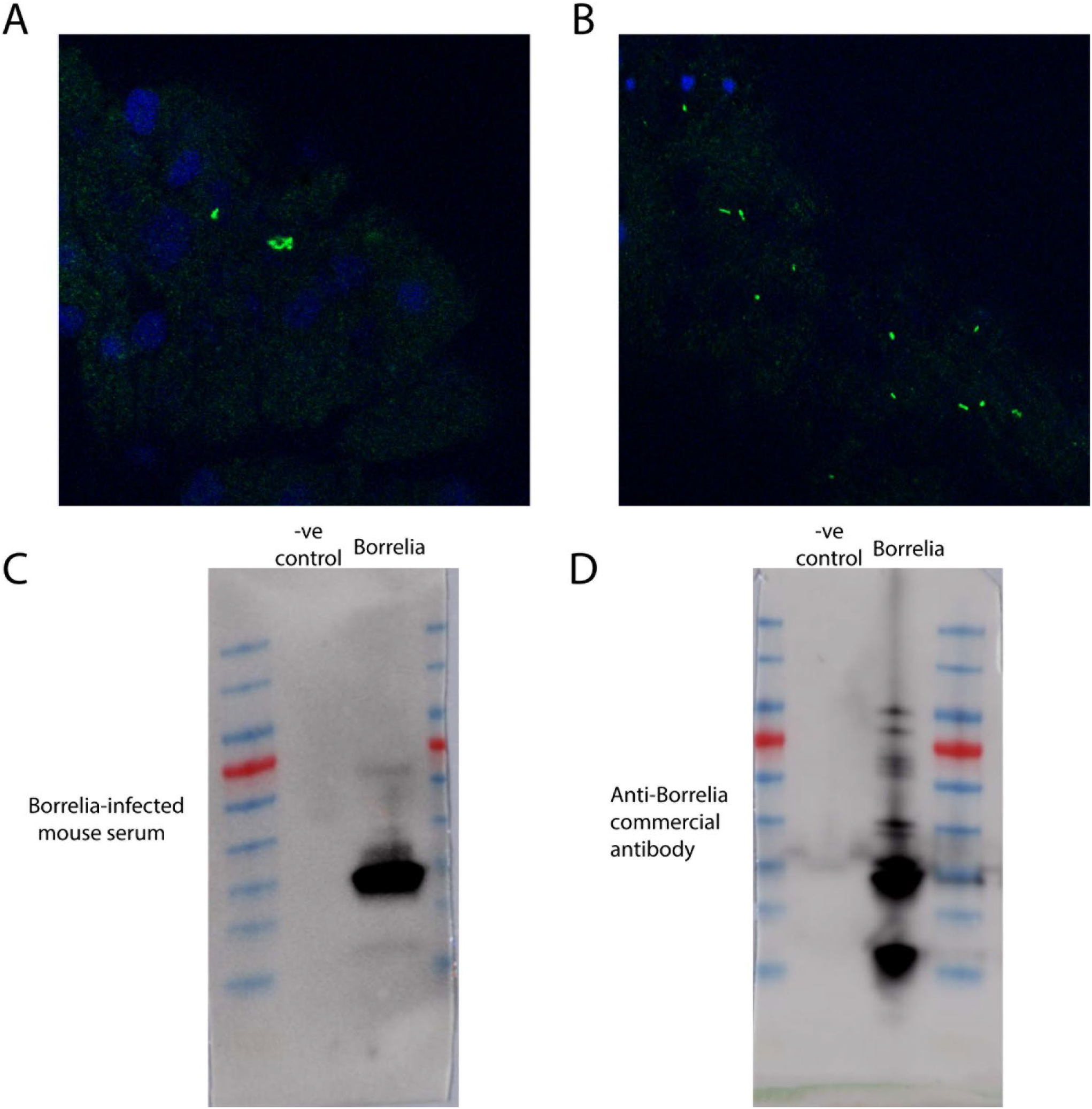
Immunofluorescence assays and western blots against *Borrelia*. (A) Immunofluorescence assay of a *Borrelia*-infected tick nymph gut utilizing anti-*Borrelia* (green) antibody and DAPI (blue). Image has been zoomed to show positive *Borrelia* signal. (B) Immunofluorescence assay of a segment of *Borrelia*-infected tick nymph gut utilizing anti-Borrelia (green) and DAPI (blue). (C) A western blot of mammalian cell lysate and cultured*Borrelia* probed with mouse serum collected post-*Borrelia* infection. (D) A western blot of mammalian cell lysate and cultured *Borrelia* probed with a commercially produced anti-*Borrelia* antibody.

## Discussion

We have detailed a general standardized methodology for studying *B. burgdorferi* transmission and acquisition on mice. Many of the conditions described herein can be altered to suit specific experimental needs.

## Author Contributions

Conceptualization, original draft preparation, review, and editing: A.S., B. L. and S.D. All authors have read and agreed to the published version of the manuscript.

